# Light affects behavioral despair involving the clock gene *Period 1*

**DOI:** 10.1101/2020.05.07.082156

**Authors:** Iwona Olejniczak, Jürgen A. Ripperger, Federica Sandrelli, Anna Schnell, Laureen Mansencal-Strittmatter, Ka Yi Hui, Andrea Brenna, Naila Ben Fredj, Urs Albrecht

## Abstract

Light at night has strong effects on physiology and behavior of mammals. It affects mood in humans, which is exploited as light therapy, and has been shown to reset the circadian clock in the suprachiasmatic nuclei (SCN). This resetting is paramount to align physiological and biochemical timing to the environmental light-dark cycle. Here we provide evidence that light affects mood-related behaviors also in mice by activating the clock gene *Period1 (Per1)* in the lateral habenula (LHb), a brain region known to modulate mood-related behaviors. A light pulse given at ZT22 to wild type mice caused profound changes of gene expression in the mesolimbic dopaminergic system including the nucleus accumbens (NAc). Sensory perception of smell and G-protein coupled receptor signaling was affected the most in this brain region. Interestingly, most of these genes were not affected in *Per1* knock-out animals, indicating that induction of *Per1* by light serves as a filter for light-mediated gene induction in the brain. Taken together we show that light affects mood-related behavior in mice at least in part via induction of *Per1* in the LHb with consequences on signaling mechanisms in the mesolimbic dopaminergic system.

## Introduction

The circadian clock has evolved from cyanobacteria to humans in response to the daily light-dark cycle (Rosbash, 2009). The internalization of the regularly recurring alternation of light and darkness allowed organisms to anticipate this change. This enabled them to prepare biochemical and physiological processes in order to optimally respond to the upcoming daily challenges and increase survival in a competitive environment. Malfunctioning or disruption of the circadian clock system results in mammals in various pathologies including obesity, cancer and neurological dysfunctions (Roenneberg and Merrow, 2016). For the maintenance of synchronicity within the mammalian body and with the environmental light-dark cycle, the suprachiasmatic nuclei (SCN) receive light information directly from intrinsically photosensitive retinal ganglion cells (ipRGCs) (Berson et al., 2002; Hattar et al., 2002; Provencio et al., 2000). This information is converted by the SCN into humoral and neuronal signals to set the phase of circadian oscillators and drive circadian rhythmic coherence in the body (Morin, 2013).

At the cellular level, the clock mechanism is relying on feedback loops involving a set of clock genes. In mammals the main feedback loop consists of two *period* (*Per1* and *Per2*) and two *cryptochrome* genes (*Cry1* and *Cry2*), whose transcription is controlled by the transcriptional activators BMAL1 and CLOCK (or NPSA2). PER and CRY form complexes with additional proteins, enter the nucleus and inhibit the activity of BMAL1/CLOCK complexes stopping their own activation. A second feedback loop involving the nuclear orphan receptors REV-ERB (α, β) and ROR (α, β, γ) regulates the expression of the *Bmal1* and *Clock* genes, whose proteins in turn regulate the *Rev-erb* and *Ror* genes (reviewed in Takahashi, 2017). Posttranslational modifications modulate the activity and stability of clock proteins and thereby contribute to and fine-tune circadian rhythm generation (Robles et al., 2017).

In order to adjust the clock to environmental signals such as light, at least one of the clock components needs to be responsive to the stimulus. This leads to a shift of clock phase in order to synchronize the organism to environmental time. Interestingly, expression of the *Per1* gene is inducible in the SCN by a nocturnal light pulse at zeitgeber time (ZT) 22 (Albrecht et al., 1997; Shigeyoshi et al., 1997). At this time point, light not only causes eventual adaptation of the mammalian circadian clock to a new time zone but is also most effective in the treatment of some forms of depression, such as seasonal affective disorder (Wirz-Justice et al., 2013). That light exerts powerful effects on mood and cognition has been documented not only in humans (Kripke et al., 1983; Lewy et al., 1982; Vandewalle et al., 2010), but also in laboratory animals (Bedrosian et al., 2011; LeGates et al., 2012; Schulz et al., 2008). The neural basis for the influence of light on mood and learning appear to be distinct retina-brain pathways involving ipRGCs that either project to the SCN to influence learning or to the perihabenular nucleus (PHb) in the thalamus to modulate mood (Fernandez et al., 2018). Furthermore, antidepressive effects of light therapy require activation of a pathway leading from the retina via the ventral lateral geniculate nucleus (vLGN)/intergeniculate leaflet (IGL) to the lateral habenula (LHb) (Huang et al., 2019).

Here we find that light induces *Per1* gene expression in the LHb and suppression of it increases immobility in the FST. Furthermore, lack of *Per1* in mice increases the number of light-induced genes in the SCN and LHb, while decreasing this number in the NAc. The profound changes in gene expression in the mesolimbic dopaminergic system are linked to sensory perception of smell and G-protein coupled receptor signaling. Our observations suggest an involvement of *Per1* in the light-mediated pathway that regulates mood-related behaviors.

## Results

### Light at ZT22 affects despair-based behavior

In order to test the effect of light on mood-related behavior in mice, we applied a 30-min polychromatic light pulse (white light) to animals that were kept in a 12-hour light/12-hour dark (12:12 LD) cycle. The light pulse was applied in the dark phase at zeitgeber time (ZT) 14 or 22, respectively, with ZT0 being lights on and ZT12 lights off. The mice were then subjected to a forced swim test (FST) at ZT6 for the next five days where their immobility time was assessed (Fig. 1A, S1A). Animals that received no light pulse (black bars) or a light pulse at ZT14 (red bars) showed comparable immobility times (Fig. 1A, S1A). In contrast, mice that received a light pulse at ZT22 (blue bars) displayed lower immobility times compared to control animals. This difference was statistically significant in females (Fig. 1A), but only showed a similar trend in males (Fig. S1A), which is consistent with previous findings in rats (Schulz et al., 2008).

**Figure 1.**
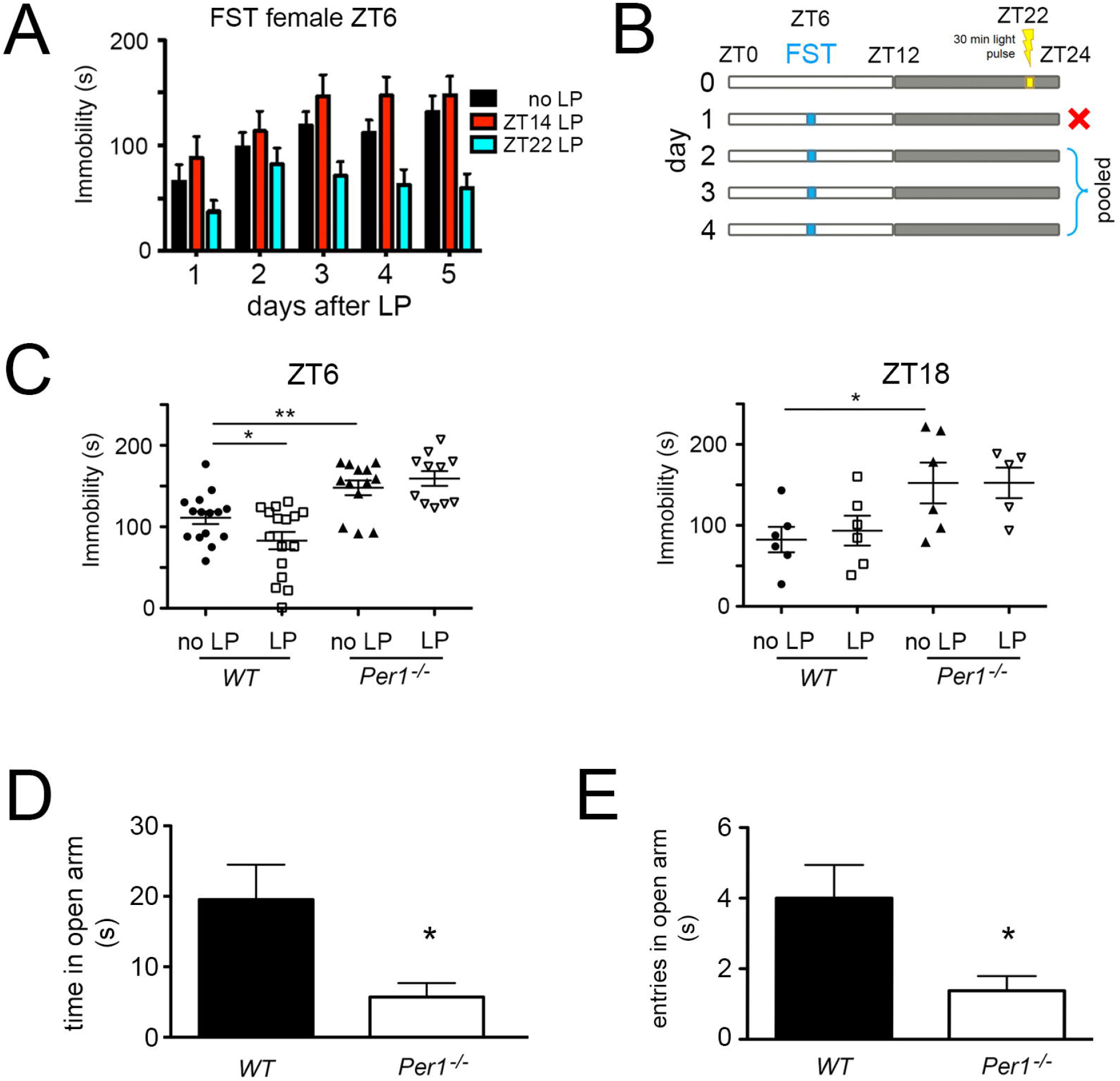
Light at ZT22 affects despair-based behavior. (**A**) Immobility time in the forced swim test (FST) of wild type female mice assessed over several days at ZT6 after no light pulse (LP) (black bars), after a LP at ZT14 (red bars), and after a LP at ZT22 (blue bars). Two-way RM ANOVA (n=13-15), ZT14 LP p=0.255, ZT22 LP p=0.014, values are means ± SEM (**B**) Light pulse treatment and assessment protocol using the FST. The first day after the light pulse the FST was performed but only the data from days 2-4 were pooled and used for further analysis. (**C**) Immobility time of wild type and *Per1*^-/-^ female mice with and without LP at ZT22 assessed at ZT6 (left panel, n=11-16) or ZT18 (right panel, n=5-6). Unpaired t-test, *p<0.05, **p<0.01, values are means ± SEM. (**D**) Time spent in the open parts of an O-maze. Unpaired t-test, n=8 for each genotype, *p<0.05, values are means ± SEM. (**E**) Entries into the open parts of an O-maze. Unpaired t-test, n=8 for each genotype, *p<0.05, values are means ± SEM.

Based on these data we performed the subsequent experiments with female mice and applied a light pulse only at ZT22. The FST was performed at ZT6 and immobility time was assessed for the four following days. The data of days 2-4 were pooled to compare immobility times between animals (Fig. 1B). Consistent with the results in Fig. 1A wild type mice showed reduced immobility time at ZT6 when they received a light pulse at ZT22 (Fig. 1C, left panel), but not when they were assessed at ZT18 (Fig. 1C, right panel). Since *Per1* is a light-inducible gene in the SCN (Albrecht et al., 1997; Shigeyoshi et al., 1997), we tested mice lacking this gene in the same paradigm. At ZT6, as well as at ZT18, *Per1* knock-out (*Per1*^-/-^) mice show increased immobility time compared to wild type animals, and a light pulse did not affect their immobility time in the FST (Fig. 1C). Hence, *Per1*^-/-^ animals showed an increase in behavioral despair, which was not decreased by light in contrast to wild type animals. Next, we used the sucrose preference test, which provides a measure for anhedonia, another characteristic of depression (decreased ability to experience pleasure; Crawley, 2000). We observed no difference between wild type and *Per1*^-/-^ mice in this experiment (Fig. S1B). To test anxiety-related behavior in the two genotypes we used the elevated O-maze test (Crawley, 2000). The *Per1*^-/-^ mice spent less time in the open area and did enter it significantly less compared to wild type control animals (Fig. 1 D, E). These results indicated that *Per1*^-/-^ animals were more anxious than controls.

### Light at ZT22 induces *Per1* in the lateral habenula

Because the LHb is involved in the regulation of the behavioral response of mice in the FST (Huang et al., 2019), we performed in situ hybridization experiments on mouse brain sections in order to evaluate, whether the *Per1* and *Per2* genes were expressed in the LHb and the medial habenula (MHb). We observed, that *Per1* was expressed in both the LHb and MHb although with a much lower amplitude than in the SCN (Fig. 2A). *Per2* mRNA was expressed in both the LHb and the MHb but to a much lower extent (Fig. 2B). Since light at ZT22 affects immobility in the FST (Fig. 1C), and induces *Per1* but not *Per2* expression in the SCN (Albrecht et al., 1997; Shigeyoshi et al., 1997) we tested the effect of a light pulse at ZT22 on expression of the *Per1* gene. We detected strong induction of *Per1* in the SCN one hour after the light pulse, which was weaker after two hours (Fig. 2C, left panel) consistent with previous findings (Albrecht et al., 1997; Shigeyoshi et al., 1997). A similar pattern of *Per1* gene induction was observed in the LHb although this was less pronounced when compared to the SCN (Fig. 2C, right panel). Interestingly, a light pulse at ZT14 did not induce *Per1* in the LHb contrary to the SCN (Fig. S2A) paralleling the absence of a light effect in the FST (Fig. 1A).

**Figure 2.**
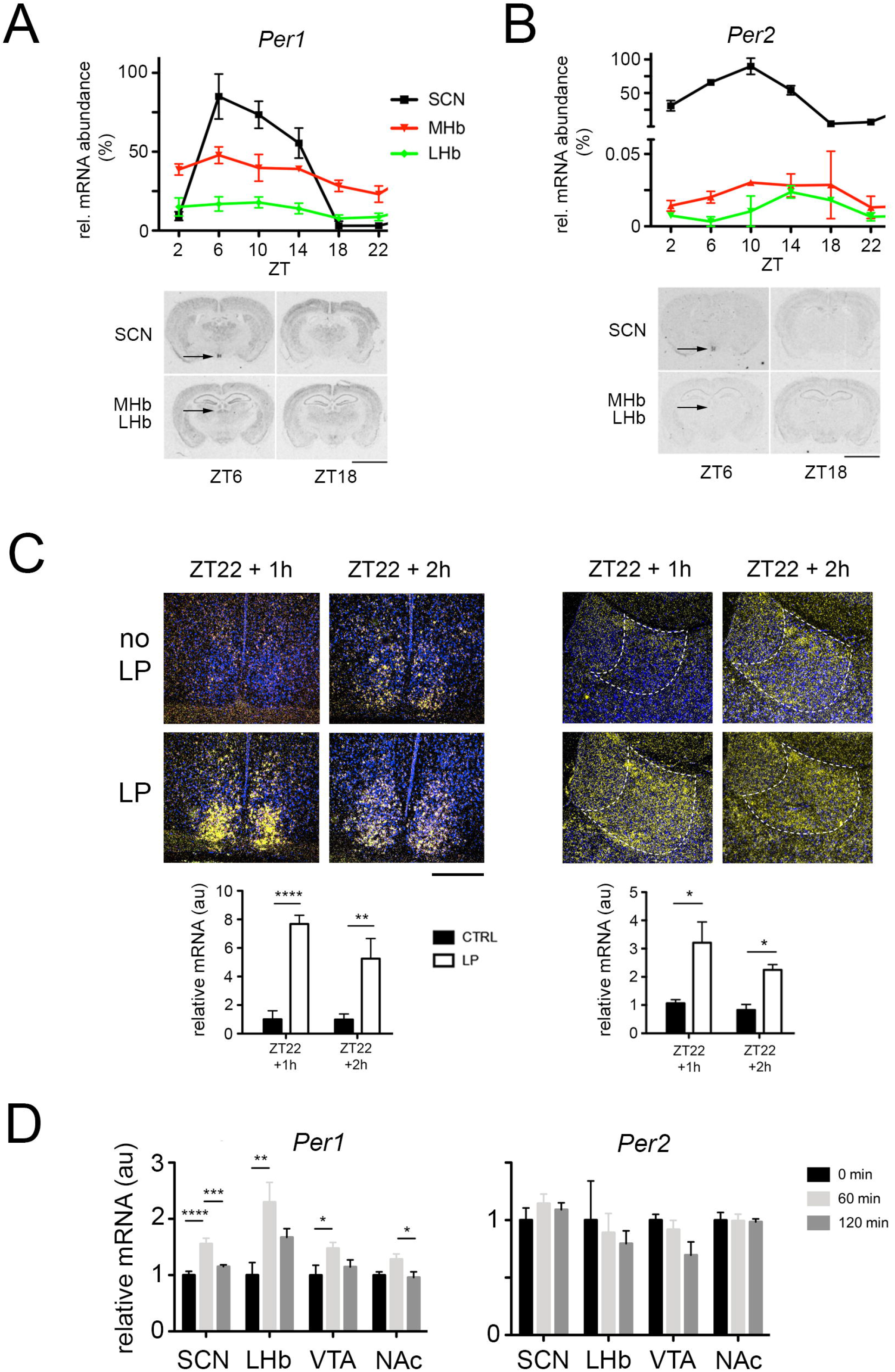
Light induces the clock gene *Per1* in the lateral habenula of mice. In situ hybridization revealing expression of *Per1* (**A**) or *Per2* (**B**) in the SCN, the medial habenula (MHb) and the lateral habenula (LHb). The top panel shows the quantification of *Per1* expression in the SCN, the MHb and the LHb, values are means ± SEM. CircWave analysis revealed circadian expression of both genes in the tissues shown (n=3, p<0.05). The bottom panel shows representative images of brain sections at ZT6 and ZT18 with expression signal (black) of *Per1* or *Per2* in the respective brain regions. Scale bar: 5 mm. (**C**) Induction of *Per1* mRNA expression after a LP at ZT22 in the SCN (left panels) and in the lateral habenula (right panels). The blue color depicts cell nuclei (Hoechst staining) and the yellow color shows radioactively labeled antisense *Per1* riboprobe hybridized to *Per1* mRNA. Below the photo-micrographs quantification is shown. Two-way ANOVA with Bonferroni multiple comparisons test, n=3, *p<0.05, **p<0.01, ****p<0.0001, values are means ± SEM. Scale bar = 200 μm. (**D**) Quantitative PCR revealed mRNA expression levels of *Per1* (left panel) and *Per2* (right panel) in the suprachiasmatic nuclei (SCN), lateral habenula (LHb), ventral tegmental area (VTA) and nucleus accumbens (NAc) after 60 min. (light grey bars) and 120 min. (dark grey bars) of a light pulse at ZT22 (0 min., black bars). *Per1* mRNA was induced in the SCN, LHb, and VTA, whereas *Per2* was not induced in any of the tissues investigated. One-way ANOVA, Tukey’s multiple comparisons test, n=12-16, *p<0.05, **p<0.01, ***p<0.001, ****p<0.0001, values are means ± SEM.

To corroborate our observations, we performed quantitative real-time PCR analysis. In order to demonstrate the accuracy of the isolation of SCN, LHb, VTA, and NAc tissue from wild type mouse brains, we used markers specific to the corresponding brain region (Fig. S2B). Mice that received a light pulse at ZT22 were compared to controls not receiving any light. We detected induction of *Per1* but not *Per2* in the SCN, the LHb and the VTA one hour after the light pulse (Fig. 2D), which was consistent with the data we obtained by using in situ hybridization.

Taken together our data suggest a role of *Per1* in the light-mediated effects on despair related behavior and that this may involve the LHb.

### Knock-down of *Per1* in the area of the lateral habenula affects despair and anxiety-related behavior

In order to challenge the hypothesis that expression of *Per1* in the LHb plays an important role in the regulation of despair based and anxiety-related behavior, we knocked down *Per1* in the LHb. To this end we tested short hairpin RNAs (shRNAs) against *Per1* in NIH 3T3 fibroblast cells (Fig. S3A) and used the variant A in the subsequent experiments. To ensure efficient delivery of the shPer1 RNA into the LHb we tested the transduction efficiency and tissue penetration of various serotypes of adeno-associated virus (AAV) expressing GFP (Fig. S3B). We injected AAV6 containing expression vectors for either a scrambled set of shRNA or a *Per1*-specific shRNA into the LHb (variant A, Fig. S3A). Examples visualizing the infection sites are shown in Fig. 3A. After recovery from the procedure the animals were tested for efficiency of *Per1* knock-down. A significant decrease of *Per1* expression was observed (Fig. 3B). After establishing the procedure, injected animals were subjected to the FST at ZT6 with shPer1 animals displaying a significantly increased immobility time compared to the scramble (scr) controls (Fig. 3C). This suggested that reduction of *Per1* expression in the area of the LHb increased despair related behavior, consistent with the observation in total *Per1* knock-out animals (Fig. 1C). We also evaluated the *Per1* knock-down animals (shPer1) in the elevated O-maze test. The time of the shPer1 animals spent in the open arm (Fig. 3D), as well as the number of entries into the open arm (Fig. 3E), displayed a trend towards lower values compared to control (scr) mice. This trend was consistent with the observations in *Per1*^-/-^ mice (Fig. 1D, E). The analysis of head dips in the O-maze showed a tendency to decreased values in the shPer1 animals (Fig. 3F) with the number of stretch-attend postures being significantly reduced in shPer1 mice compared to scr controls (Fig. 3G). Overall these results indicate that *Per1* expression in the habenula counterbalances despair-based and anxiety-related behaviors.

**Figure 3.**
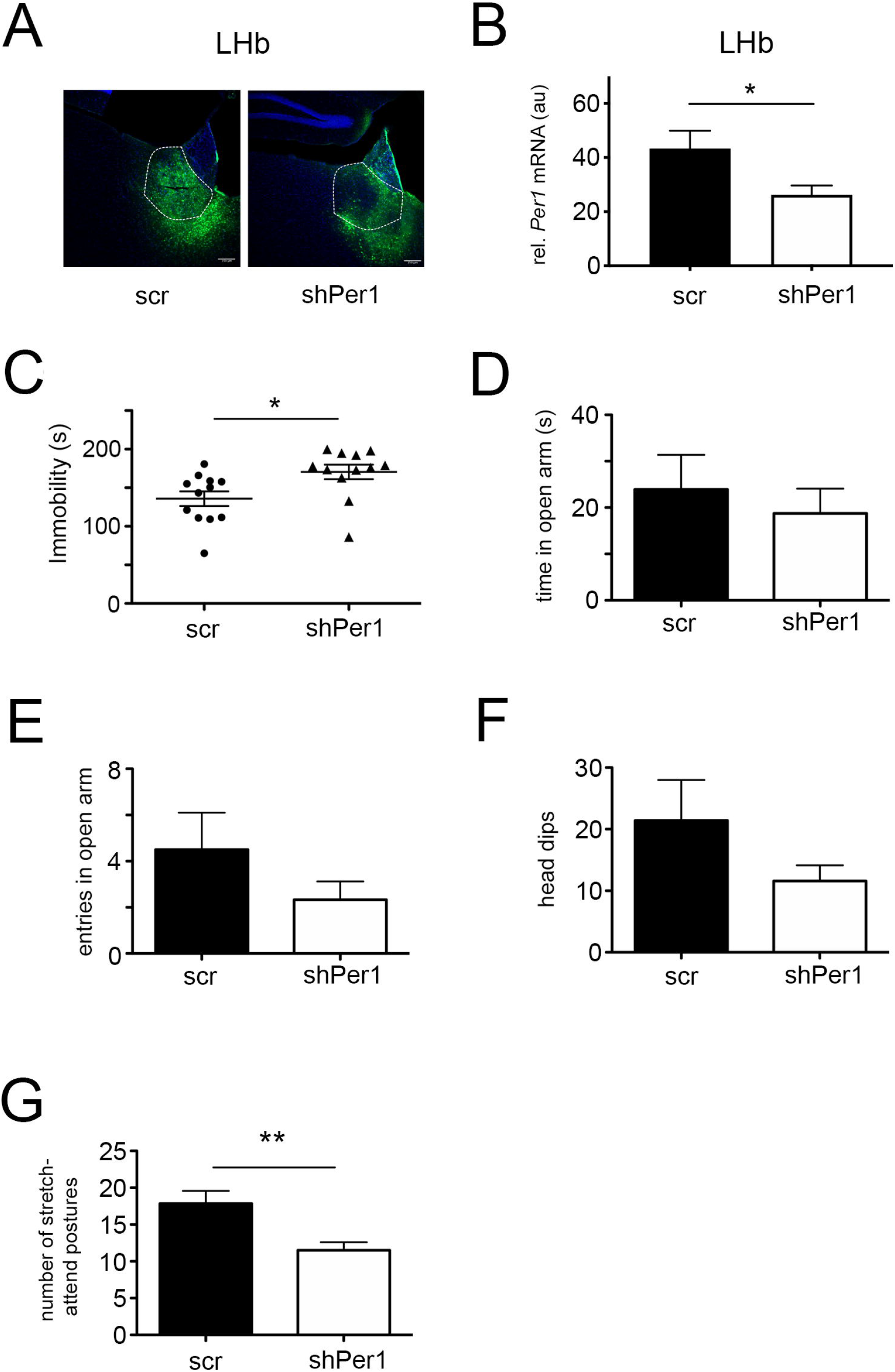
Knock-down of *Per1* in the area of the lateral habenula affects despair and anxiety-related behavior. (**A**) Injection sites of adeno-associated virus (AAV6) into the area of the lateral habenula expressing a scrambled (scr) or shRNA against *Per1* (shPer1). Scale bar: 150μm. (**B**) Quantification of knock-down efficiency in the LHb. *Per1* expression is significantly reduced by shPer1 compared to scr. Unpaired t-test, n=6, *p<0.05. (**C**) Immobility time in the FST of female mice at ZT6 injected with AAV expressing scr or shPer1. Unpaired t-test, n=12, *p<0.05. (**D**) Time in open arm of an O-maze at ZT6. Unpaired t-test, n=12, p>0.05. (**E**) Number of entries in open arm of an O-maze at ZT6. Unpaired t-test, n=12, p>0.05. (**F**) Head dips of mice in the O-maze at ZT6. Unpaired t-test, n=12, p>0.05. (**G**) Number of stretch-attend postures of mice in the O-maze at ZT6. Unpaired t-test, n=12, **p<0.01, values in all experiments are means ± SEM.

### Light induction of PER1 protein shapes the transcriptome in the brain in a regional manner

Presence or absence of the *Per1* gene changes behavior of mice in the FST. In order to relate this behavioral effect to alterations in the transcriptome, we performed RNA sequencing experiments. Because we were interested in the downstream impact of *Per1*, we first studied the amount of PER1 protein after the light pulse at ZT22 in the SCN and the LHb. Previous studies showed, that after a light pulse at ZT22 murine PER1 protein in the SCN was increased 10 hours later (ZT8) when compared to mice that received no light (Yan and Silver, 2004). We confirmed this observation in the SCN by Western blot and found a similar increase of PER1 protein in the LHb (Fig. 4A). This suggested that *Per1* is light-inducible in the LHb.

**Figure 4.**
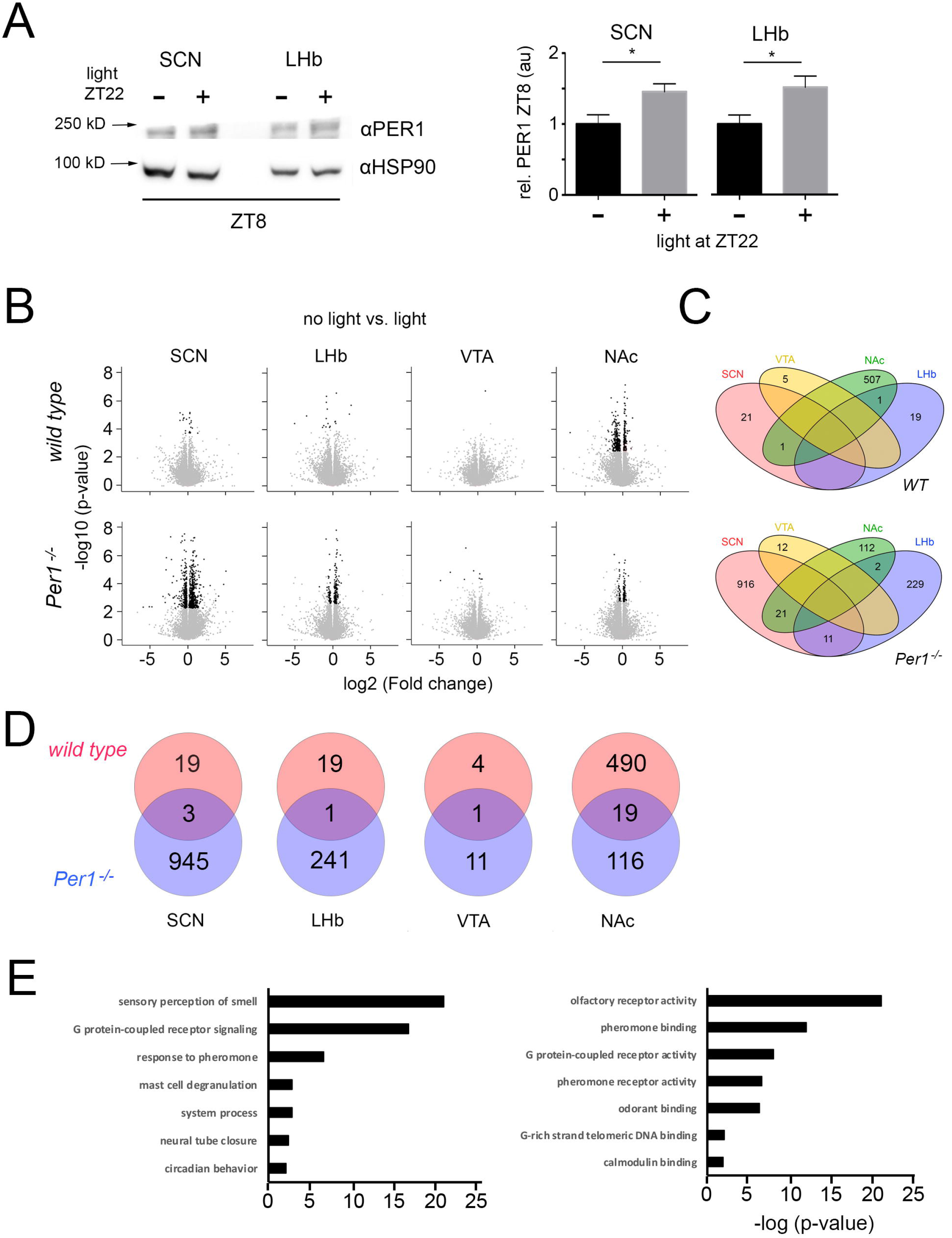
Light induction of PER1 protein at ZT8 and effects on the transcriptome in the brain. (**A**) Expression of PER1 protein at ZT8 in the SCN and the LHb after a light pulse at ZT22 compared to no light pulse. The left panel depicts an example of a Western blot and the right panel shows the quantification of three independent experiments. Unpaired t-test, n=3, *p<0.05, values are means ± SEM. (**B**) Volcano plots depicting the changes of gene expression in response to a light pulse at ZT22 in the indicated tissues of wild type and *Per1*^-/-^ mice. The RNA sequencing was perfomed 10 hours after the light pulse (ZT8). Black dots indicate significant changes, n=6. (**C**) Venn-diagrams illustrating the overlap of gene expression changes in the four nuclei. Top comparison in wild type animals, bottom comparison in *Per1*^-/-^ mice. Numbers indicate the amount of genes affected by the light pulse. For the gene names see table 1. (**D**) Venn-diagram comparing wild type and *Per1*^-/-^ mice in a single nucleus. Numbers indicate number of genes affected by the light pulse. Genes are listed in the tables; SCN: wt (table 2), *Per1*^-/-^ (table 3); LHb: wt (table 4), *Per1*^-/-^ (table 5); VTA: wt (table 6), *Per1*^-/-^ (table 7); NAc: wt (table 8), *Per1* (table 9). (**E**) TopGO analysis of biological processes (left) and molecular functions (right) in the NAc of wild type animals.

Because PER1 protein presence and hence its biological function is increased at ZT8 after a light pulse at ZT22 compared to controls, we collected brain tissues at ZT8. We chose the SCN as reference tissue and the LHb, VTA and NAc as tissues known to be involved in the regulation of behaviors related to mood and despair as part of the mesolimbic dopaminergic system (Huang et al., 2019; Nestler and Carlezon, 2006). The tissues from wild type and *Per1*^-/-^ mice receiving no light or light at ZT22 were isolated at ZT8. We validated the specificity of the tissues isolated using marker genes such as lrs4 (SCN), Gpr151 (LHb), Tacr3 (VTA) and Tac1 (NAc) (Fig. S4) that were determined from the Allan brain atlas and normalized according to Fonseca Costa et al., 2017. RNA sequencing was performed and the volcano plots comparing gene expression under no light versus light at ZT22 for the various tissues and genotypes are shown in Fig. 4B. In wild type mice the SCN and LHb showed several genes being induced 10 hours after the light pulse (black dots) with the VTA displaying a much lower number and the NAc showing a massive number of induced genes. In contrast, the change in gene induction is opposite in the *Per1*^-/-^ mice. A large number of genes induced by light were observed in the SCN and LHb with low numbers in the VTA and the NAc. These results are consistent with the role of PER1 as a repressor in the regulation of transcription (Takahashi, 2017). Since the flow of information driven by the light signal goes from the LHb via the VTA to the NAc (Huang et al., 2019) the gene expression in the NAc has an opposite dynamic in presence or absence of PER1 (Fig. 4B).

Next, we compared the genes altered in the different brain regions and genotypes (Fig. 4C). First and most strikingly, all of the genes influenced by light in the VTA, irrespective of the genotype, were specific for this region only, with no *Per1* mRNA detectable. *Per1* was not detectable, because were looking for downstream effects of *Per1* gene induction 10 hours after the light pulse when PER1 protein is high (Fig. 4A) and its mRNA was already gone. Second, with the exception of two genes, all the light-induced genes in wild type mice were brain region specific. Conversely, in the absence of *Per1* many genes in the NAc and LHb were detected in the SCN as well (21 and 11, respectively). This indicated that many signaling pathways that respond to light and at ZT8 modulate targets common to several brain regions are suppressed by PER1. Hence, PER1 appeared to contribute to brain region specific shaping of molecular responses to light.

We evaluated tissue-specific differences between the genotypes by comparing for each brain region of interest the expression between wild type and *Per1*^-/-^ mice (see tables 1-8). Most of the genes induced by light in wild type animals appeared to be directly or indirectly regulated by *Per1*, because only a small fraction of those genes were common to the light-induced genes found in *Per1*^-/-^ mice (Fig. 4D). Furthermore, we found in wild type the highest number of genes to be induced in the NAc (490) and the lowest number in the VTA (4). These differences in numbers were probably due to the fact that the signal progressing from the LHb to the NAc is a dynamic process and that ZT8 was the optimal time point to detect changes in the NAc, because PER1 protein levels were high at that time point. Interestingly, the number of light-induced genes was higher in all brain regions of *Per1*^-/-^ mice, except for the NAc (Fig. 4D). This observation highlights the repressive function of PER1 on pathways which inhibit gene expression in the NAc at ZT8.

We were most interested in the 490 genes that were induced by light at ZT8 only in the NAc of wild type mice and appeared to be depending on *Per1* gene expression. Some of these genes are likely involved in the light-mediated effects on behavioral despair that we observed (Fig. 1C). We performed a TopGo analysis of this gene set in order to relate these genes to biological processes and molecular functions (Fig. 4E). Interestingly, G protein-coupled receptor signaling and sensory perception of smell were the highest-ranking biological processes. This finding was mirrored in the molecular function analysis where G protein-coupled receptor activity and olfactory receptor activity ranked highest (Fig. 4E). This result is in line with previous observations that showed depression-like behaviors in rats after olfactory bulb removal (Leonard, 1984). Interestingly, also patients with major depression exhibited olfactory deficits (Pause et al., 2001).

## Discussion

A growing body of evidence supports the notion that light therapy can be efficient in the treatment of individuals with seasonal and non-seasonal depression (Golden et al., 2005; Kripke, 1998; Lam et al., 2016; Lieverse et al., 2011; Rosenthal et al., 1984; Sit et al., 2018; Wirz-Justice et al., 2011), while light deprivation can increase depressive-like behavior in various species (Gonzalez and Aston-Jones, 2008; Lau et al., 2011; Monje et al., 2011; Wilson, 2002). However, many unresolved questions remain regarding how light therapy can produce its beneficial effects. In this study, we offer evidence that light-mediated induction of the clock gene *Per1* in the LHb is involved in the anti-depressant effects of light therapy.

We studied the effect of light on behavioral despair in mice and uncovered a significant contribution of the clock gene *Per1* in this process. Similar to the observations in humans, where light exerts powerful effects on mood and cognition (Kripke et al., 1983; Lewy et al., 1982; Vandewalle et al., 2010), we observed that light at night affected mice in a comparable way (Fig. 1A), although mice are nocturnal and not diurnal. We found that light affected behavior in a despair-based paradigm when given in the late part of the dark phase (ZT22). On the other hand, we did not observe any change in this behavior when light was given in the early period of the dark phase (ZT14), as has been reported for rats (Schulz et al., 2008). This is very similar to light treatment being more efficient in the early morning than evening for patients with seasonal affective disorder (SAD) (reviewed in Wirz-Justice et al., 2013). Furthermore, we found that light treatment at ZT22 appeared to be more effective in female than male mice (Fig. 1A, S1A). However, in humans such a sex-related discrepancy wasn’t observed in patients treated for SAD with light (Knapen et al., 2014; Leibenluft et al., 1995). Although the incidence of depression seems to be higher in females compared to males (Rubinow and Schmidt, 2019).

Interestingly, we could see the light effect when we assessed animals at ZT6 but not at ZT18. This may indicate that either the light-dark transition at ZT12 may have an impact on the behavioral outcome, or that the circadian clock modulates the amount of immobility in the FST. The latter is more likely, because without any light pulse immobility at ZT18 is lower than at ZT6 in the FST (Hampp et al., 2008, Fig. 1C). Consistently, the lack of *Per1* not only resulted in the absence of a light response but also in an increased immobility time (Fig. 1C). Furthermore, it has been shown that the LHb was not activated by a dark-light transition (Huang et al., 2019).

Mood-related behavior is a complex trait. Therefore, in addition to despair, we tested sucrose preference and behavior in the elevated O-maze, which relate to anhedonia and anxiety, respectively. The results indicated that wild type and *Per1*^-/-^ mice are similar in their ability to experience pleasure (tasting of sucrose) (Fig. S1B), but *Per1*^-/-^ mice appeared to be significantly more anxious (Fig. 1D, E). Hence, *Per1* is likely involved in the despair and anxiety aspects of mood-related behaviors. This observation is extended by previous ones where *Per1* knock-out mice showed an inability to sensitize to cocaine and experience reward (Abarca et al., 2002). Altogether, these results support the notion that *Per1* participates in the regulation of reward processes to modulate mood-related behaviors.

In mammals, light at night can induce gene expression in the SCN (Rusak et al., 1990). The clock gene *Per1* is among these light inducible genes (Albrecht et al., 1997; Shigeyoshi et al., 1997). Since the ipRGCs not only project to the SCN, but also to other brain regions (Hattar et al., 2006), we investigated the habenula (Huang et al., 2019) for light-inducible *Per1* gene expression. We observed that *Per1* was expressed in an oscillating fashion in the lateral (LHb) and medial habenula (MHb) (Fig. 2A), while *Per2* was expressed at a much lower level in these structures (Fig. 2B). *Per1* gene expression was induced by light in the LHb when applied at ZT22 (Fig. 2C), but not when applied at ZT14 (Fig. S2A). These results indicated a time-specific inducibility of *Per1* in the LHb, which mirrored the effect of light at ZT22 observed in the FST as immobility (Fig. 1A). Our findings suggest that light induction of *Per1* in the LHb and probably in other brain regions played an important role (Fig. 2D). Of note is that *Per2* was most likely not involved, because its gene induction by light at ZT22 was minimal or absent (Fig. 2D). Interestingly, light at ZT22 elicits phase-advances, which was abrogated in mice lacking *Per1* but not *Per2* (Albrecht et al., 2001). Hence lack of phase advance and increased immobility in the FST of *Per1* knock-out mice inversely correlated with the observation in humans in which advance of sleep phase had a positive effect on depression (Wirz-Justice et al., 2013). Taken together, these results suggested that the amount of *Per1* gene expression in the SCN and the LHb correlate with despair-based behavior as observed in the FST (Fig. 1C).

In order to challenge the hypothesis that the amount of *Per1* gene expression in the LHb was relevant for the level of immobility time in the FST, we knocked down *Per1* by injecting AAVs expressing shRNA against *Per1* into the area of the LHb (Fig. 3A, B). We observed a similar increase of immobility in the FST as for the *Per1* knock-out animals (Fig. 1C), which suggested a functional relationship between *Per1* expression in the LHb and the immobility time in the FST. The importance of the LHb area in mood regulation has been described previously (Namboodiri et al., 2016). Light modulated LHb activity via M4-type melanopsin-expressing retinal ganglion cells (RGCs) thereby regulating depressive-like behaviors (Huang et al., 2019). Our observations support these findings, although we do not know whether *Per1* is induced in the LHb via the M4-type melanopsin-expressing RGCs. Interestingly, another study described light effects via intrinsically photosensitive RGCs on the SCN and the perihabenula (PHb), regulating hippocampal learning or mood, respectively (Fernandez et al., 2018). Although our experimental set up was different from that study, our findings are not contradictory, because we cannot exclude an involvement of the PHb in our study. The knock-down of *Per1* in the area of the LHb had a less pronounced effect on anxiety (Fig. 3D-G) compared to the *Per1* knock-out (Fig. 1D, E). Although the time spent in the open arm and the entries in the open arm of the shPer1 mice were tendentially similar to those of *Per1*^-/-^ animals, they were not significant. This was probably due to the fact that the knock-down of *Per1* did not block all *Per1* mRNA (Fig. 3B), and therefore the effects on the behavior of mice in the elevated O-maze were less pronounced. In addition, *Per1*^-/-^ mice lack the gene in all cells of the body. Hence, we can not exclude that other brain regions or body tissues may have contributed to the phenotype as well. Overall, our results and data by others support the notion, that the area of the LHb and expression of *Per1* are involved in the light mediated effects on mood-related behavior.

The molecular mechanisms through which light elicits its beneficial effects on mood-related behaviors are poorly understood. The observation that light induces the expression of the *Per1* gene in the area of the LHb provides an opportunity to get a first glimpse at potential molecular pathways that are initiated by the activation of *Per1*. In order to identify targets of PER1 that are involved in the behavior we observed in the FST, we performed RNA sequencing experiments. Since clock genes regulate behavior in the FST partly via the mesolimbic dopaminergic system (Chung et al., 2014; De Bundel et al., 2013; Hampp et al., 2008; Roybal et al., 2007) we used tissue from the LHb, VTA, and NAc with SCN tissue as control. The tissues were isolated at ZT8, because light-induced PER1 protein was highest at that time in the SCN (Yan and Silver, 2004) and LHb (Fig. 4A) and very close to our behavioral assessment in the FST at ZT6. We observed that light-induced genes were mostly specific to a particular tissue with virtually no overlap with other tissues assessed. In contrast, lack of *Per1* increased the overlap of induced genes between the tissues (Fig. 4C), suggesting a role of PER1 in suppressing common light-responsive genes. This evidence is consistent with the known role of PER1 as a suppressor of clock and tumor-related genes (Li et al., 2016; Takahashi, 2017). Interestingly, this can only be observed in the SCN, LHb and VTA, but not the NAc (Fig. 4D). This could be due to indirect regulation of genes in the NAc by PER1 (e.g. suppression of suppressors specific to the NAc) and/or neurotransmitter related gene regulation in the NAc by the VTA or other brain regions. Remarkably, the VTA showed the fewest number of genes induced by light. The reason for this may be the time of assessment, because at ZT8, most of the light-induced genes in the VTA may have already been silenced again. This highlights the highly dynamic nature of light-induced effects on behavior from initial gene induction, activation of signaling pathways to neuronal communication and neurotransmitter release. In order to understand this process better, a dynamic assessment of the transcriptome at several time points (e.g. every two hours after the light pulse) would be necessary. The dynamic temporal change of the transcriptome in the LHb, VTA and NAc after the light pulse may then provide insights into the relationships between the various nuclei to translate the initial light signal into a systemic change that affects mood-related behaviors. Nevertheless, our analysis of the light-induced transcriptomic change at ZT8 revealed significant contributions of genes involved in the sensory perception of smell/olfactory receptor activity and G protein-coupled receptor signaling/activity. This parallels previous findings that described depression-like behaviors in rats after olfactory bulb removal (Leonard, 1984) and patients with major depression displaying olfactory deficits (Pause et al., 2001). However, it remains elusive how the olfactory system and depression are mechanistically related.

Integrated genome-wide association and hypothalamic eQTL studies indicated a link between *Per1* and coping behavior in humans (Ponsuksili et al., 2015). Furthermore, a single nucleotide polymorphism in the human *Per1* promoter, as well as animal experimentation, revealed a role of *Per1* in psychosocial stress-induced alcohol drinking (Dong et al., 2011). These studies are consistent with our finding, that *Per1* was involved in the regulation of behavioral despair and anxiety, two aspects of depression. Interestingly, the profiles of *Per1* and *Rev-erbα* were advanced in patients in the manic phase compared with those in the depression phase (Novakova et al., 2015). This correlates with the light-induced phase-shifting properties of *Per1* when light was applied at ZT22 and eliciting a phase-advance in activity and gene expression rhythms (Albrecht et al., 1997; Albrecht et al., 2001; Shigeyoshi et al., 1997). Taken together, our study provides evidence that the benefit of light on mood-related behaviors involves the clock gene *Per1* and that induction of this gene in the area of the LHb plays an important role.

| **Key Resources Table** |  |  |  |  |
| --- | --- | --- | --- | --- |
| Reagent type (species) or resource | Designation | Source or reference | Identifiers | Additional information |
| Cell line (M. musculus) | NIH/3T3 | ATCC | Cat. #: ATCC® CRL-1658™ | Immortalized Mouse fibroblast cells |
| Antibody | anti-PER1 (Rabbit polyclonal) | Created by J.Ripperger |  | 1:1000 (WB) |
| Antibody | anti-HSP90 $\alpha/\beta$ (mouse, monoclonal) | Santa Cruz Biotechnology | Cat# : sc-13115<br>RRID:AB_627758 | 1:1000 (WB) |
| Antibody | anti-GFP (Rabbit polyclonal) | Abcam | Cat. #: ab6556<br>RRID: AB_305564 | 1:500 (IF) |
| Antibody | Alexa Fluor 488-AffiniPure Donkey Anti-rabbit IgG (H+L) (Donkey polyclonal) | Jackson ImmunoResearch | Cat. #: 711-545-152<br>Lot # 132876<br>RRID: AB_2313584 | 1:500 (IF) |
| Antibody | anti-mouse IgG – peroxidase, produced in rabbit | Sigma-Aldrich | Cat # : A9044<br>RRID: AB_258431 | 1:10000(WB) |
| Antibody | anti-rabbit IgG – peroxidase, produced in goat | Sigma-Aldrich | Cat# : A9169<br>RRID: AB_258434 | 1:10000(WB) |
| Chemical compound | 2% Lidocain HCl | Bichsel | Zul Nr. 53570089 |  |
| Chemical compound | Rimadyl (Carprofenum 50mg/ml) | Zoetis | Swissmedic: 54375018 |  |
| Chemical compound | RTI-13951-33 hydrochloride | MedChemExpress | Cat. # HY-112612A |  |
| Chemical compound | cOmplete™, EDTA-free Protease Inhibitor Cocktail | Roche Diagnostics GmbH | Cat. # 06538282001 |  |
| Commercial assay, kit | RNeasy® Micro Kit | Qiagen | Cat. # 74004 |  |
| Commercial assay, kit | SuperScript™ II Reverse Transcriptase | Invitrogen | Cat. # 18064022 |  |
| Commercial assay, kit | KAPA PROBE® FAST qPCR Master mix | Kapa biosystems | Cat. # KR0397 |  |
| Commercial assay, kit | Lipofectamine™ 2000 Transfection Reagent | Thermo Scientific | Cat. # 11668019 |  |
| Commercial assay, kit | Pierce™ Rapid Gold BCA Protein Assay Kit | Thermo Scientific | Cat. # A53225 |  |
| Commercial assay, kit | Pierce® ECL Western Blotting Substrate | Thermo Scientific | Cat. # 32106 |  |
| Genetic reagent (M. musculus) | mPer1Brdm1 | Zheng et al., 2001 |  | PMID: 11389837 |
| other | DAPI | Thermo fisher | Cat. #: D3571<br>RRID: AB_2307445 | (1 µg/mL) |
| Recombinant DNA reagent | Per1 Mouse shRNA Plasmid (Locus ID 18626) | Origene | CAT#: TL501619 | Mouse, 4 unique 29mer shRNA constructs in lentiviral GFP vector |
| Recombinant DNA reagent | Scrabled shRNA Plasmid | Origene | CAT#: TR30021 | Non-effective 29-mer Scrambled shRNA Cassette in pGFP-C-shLenti Vector, 5 ug |
| Recombinant DNA reagent | Adeno-associated virus (AAV) | Viral Vector Facility (VVF) of the Zurich Neuroscience Centre (ZNZ) | v77<br>vIO3-6<br>v344 |  |
| Recombinant DNA reagent | Primers and TaqMan probes | Complete list provided in the paper |  |  |
| Software | Leica application Suite Advanced Fluorescence | Leica | Version 2.7.3.9723 |  |
| Software | Prism | GraphPad | Version 7.0d |  |
| Software | Quantity one | Biorad | Version 4.6.2 |  |
| Software | ImageJ | ImageJ | Version 1.51n |  |
| Software | Circ Wave | Roelof A. Hut | Version 1.4 |  |

## Materials and Methods

### Animals and housing

Animals were kept in 12h light/12h dark (12:12 LD) cycle with food and water *ad libitum*. Timing of experiments is expressed as zeitgeber time (ZT, ZT0 lights on, ZT12 lights off). Up to 4 animals were housed in plastic cages (268mm long x 215mm×141 mm high; Techniplast Makrolon type 2 1264C-001) covered with a stainless-steel wire lid (Techniplast 1264C-116) and a filter top (1264C-400SC; 1264C-420R). Unless otherwise stated, both male and female mice were used in experiments. Housing and experimental procedures were performed in accordance with the guidelines of the Schweizer Tierschutzgesetz and the declaration of Helsinki. The state veterinarian of the Canton of Fribourg approved the protocol (2015-33-FR). *Per1^Brdm1^* total knock out animals (Zheng et al., 2001) PMID: 11389837 were used.

### Light pulse experiment

Mice aged 3 – 6 months were exposed to polychromatic light at 500 lux in their home cage for 30 min at ZT14 or ZT22 (until ZT22.5). The control group was subjected to the same handling (cage movement, presence of the experimenter) in the dark but receiving no light pulse.

### Stereotaxic injections

The following adeno-associated viruses (AAV) were provided by the Viral Vector Facility (VVF) of the Neuroscience Center Zürich (ZNZ): for serotype testing - v77: ssAAV-X/2-hCMV-chI-EGFP-WPRE-SV40p(A) where X indicates serotype between: 1, 2, 5, 6, 8, 9, DJ; for *Per1* in vivo knock down - RNA interference vector (AAV6) containing the sequence of microRNA (shRNA A: 5’-CAGTGTAGCTTCAGCTCCACCATCGTCCA-3’) supplied by the experimenters (vIO3-6: ssAAV-6/2-hSyn1-chI[1x(murine shPer1)]-EGFP-WPRE-SV40p(A)) and the scramble control (v344: ssAAV-6/2-shortCAG-chI[1x(shNS)]-EGFP-WPRE-SV40p(A)). AVV vectors and plasmids required for their synthesis are available on the VVF website (https://www.vvf.uzh.ch/en.html).

2-months old mice were selected for the stereotaxic injections. Under isolfurane/oxygen anasthesia (2.0% isoflurane v/v, 1 l O_2_/min, induction: 5.0% v/v, 1.0l O_2_/min) the animal’s head was fixed in the stereotaxic instrument (Kopf instruments) and fitted with an anesthesia inhalation mask (Stoelting). The depth of anesthesia was periodically monitored with the toe-pinch reflex. Bregma was used as reference point for positioning the skull and coordinates (position 0). The brain was accessed with craniotomy. The injection (200 nl at 40 nl/min) at LHb-specific coordinates (AP = −1.75mm, ML = +/-0.9mm, DV = 2.7mm, inward angle +/-10^0^) was performed with a pulled glass pipette (Drummond, 10μl glass micropipette, Cat number: 5-000-1001-X10) attached to a hydraulic manipulator (Narishige: MO-10 one-axis oil hydraulic micromanipulator). The micropipette was risen by 0.1mm and kept in place for 5 min to prevent upward spread of the virus. For analgesia 2% lidocain was applied locally and 5 mg/kg of caprofen was injected subcutaneously on the day of the surgery and during the two following days. Animals that underwent the surgery were left in their home cage for 14 days to recover.

### Behavioral testing

To reduce stress and burden for the animals gentle handling methods were applied wherever possible, including cup and tunnel handling (Ghosal et al., 2015; Hurst and West, 2010). For behavioral testing, mice were caged individually at least 4 days before the experiment.

*Forced swim test (FST)* was performed according to established protocols (Crawley, 2000) at an experiment-specific zeitgeber time. A transparent cylinder 35 cm high with a diameter of 25 cm was filled with water up to 20 cm high. The temperature of the water was kept at 25 +/- 1 °C. During a 6-minute session the animal was placed in the cylinder and allowed to swim freely while the experimenter left the room to minimize disturbance. The session was recorded horizontally with a video camera and the video was scored manually. During one session, two animals were tested at the same time in neighboring cylinders separated by a thin white wall to obstruct the view. The test result was expressed as immobility time in seconds recorded during the last 4 minutes of the test. Small movements of paws and tail meant to stabilize the animal’s position while floating were not considered as mobility. Scoring was performed blinded with the assessor not knowing the genotype or treatment of the animals.

Anxiety was measured using *elevated O-maze* test at ZT6. Mice were placed on an elevated ring platform (40 cm Ø), divided into four parts, equal in size, two exposed, and two flanked by 15 cm high walls. Each animal underwent a single 10-minute session which was recorded by video camera from above. Time in the exposed part, entries into the exposed portion (full body), head dips and stretch-attend postures were counted manually. Scoring was performed blinded with the assessor not knowing the genotype or treatment of the animals.

### Sucrose preference test

Sucrose (1, 2.5, and 5% w/v) solution intake was measured in a two-bottle free choice test (sucrose against water) (Spanagel et al., 2005). After weighing, mice were placed into individual cages at least three days before the start of the experiment. One day before the beginning of the experiment mice were habituated to the presence of two bottles in the cage both containing water. After 12 hours the consumed water for each bottle was measured and the position of the bottles was inverted. At the start of the test, one water bottle and one bottle containing a sucrose solution (1, 2.5 or 5% w/v) were introduced to each cage. Each sucrose concentration was present for three days). The amount of consumed water or sucrose solution in 24 hours was measured with exchanging the position of the bottles every 12 hours. Sucrose preference was calculated by dividing the amount of sucrose consumed by the total amount of consumed solutions (water and sucrose). Mice were weighed every day.

### Brain region isolation

A freshly isolated brain was cut on a brain matrix. The regions of interest were cut out of the corresponding brain slice and immediately frozen in liquid nitrogen. Tissue was stored in −80 °C until further analysis.

### Immunostaining

Brain tissue was washed with 0.9% NaCl, 10KU/l Heparin and subsequently fixed with 4% PFA by cardiovascular perfusion. The tissue was cryoprotected with 30% sucrose/1xPBS (137 mM NaCl, 7.97 mM Na_2_HPO_4_ × 12 H_2_O, 2.68 mM KCl, 1.47 mM KH_2_PO_4_). Coronal sectioning was performed on a cryostat (Microm, Thermo Scientific, HM550) at 30 μm section thickness. Pre-selected slices were washed for 5 min once in 1xTBS (0.1 M Tris/0.15 M NaCl) and twice in 2xSSC (0.3 M NaCl/0.03 M tri-Na-citrate pH 7). Antigen retrieval comprised of 50 min incubation in 65 °C in 2xSCC/50% Formamide. Subsequently, the slices were washed 2 times, 5 minutes in 2xSSC, 3 times in 1xTBS and blocked for 1 hour at 25 °C (10%FBS/1xTBS/0,1%Triton). Primary antibody (anti-GFP 1:500, Abcam, ab6556) was diluted in 1%FBS/1xTBS/0.1% Triton and incubated for 18h at 4 °C. Next the sections were washed 3 times with 1xTBS. Secondary antibody (Alexa Fluor^®^ 488 AffiniPure Donkey Anti-Rabbit IgG (H+L), Jackson Immunoresearch, ref: 711-545-152, 1:500 in 1%FBS/1xTBS/0.1% Triton) was added for a 3-hour incubation at 25 °C, followed by 3 washes in 1xTBS, 10 min DAPI staining (1:5000 in PBS; Roche) and 2 washes with 1xTBS. The slices were mounted on slides (SlowFade™ antifade reagent, Invitrogen) and visualized with a confocal microscope.

Confocal pictures were taken using a Leica TCS SP5 microscope with a 10x objective. Resolution of the image was set to 1024×1024 pixels, scanning speed to 200 Hz with 10% laser power. Images were captured using Leica LAS (2.7.3) software and processed using ImageJ (1.51n).

### Quantitative polymerase chain reaction (qPCR) analysis

RNA from tissue fragments was isolated with RNeasy® Micro Kit (Qiagen) according to manufacturer’s instructions. Reversed transcription was performed using SuperScript™ II Reverse Transcriptase (Invitrogen) according to the manufacturers protocol (250 - 1000 ng RNA per reaction). After reversed transcription, samples were diluted 10 times. 5μl of the diluted sample was added to 7.5μl of KAPA mix (KAPA PROBE^®^ FAST qPCR Master) and 2.5 μl of primers/probe mix (listed in table 10) (150 nM of primers, 33.3 nM TaqMan^®^ probe). Samples were analyzed using the Rotor-Gene qPCR machine (Corbett research, RG-6000). For the primers used see Table 1.

### Transcriptome analysis

Female mice (4.5 months of age) were sacrificed at ZT8 (10 h after a 30 min light pulse at ZT22) and brain regions were isolated as described above. For RNA isolation, two animals were pooled for each sample (total of 12 animals analyzed in 6 separate samples) for each phenotype and treatment group (wild-type, wild-type plus light pulse, *Per1*^-/-^, *Per1*^-/-^ plus light pulse) to match a total of 24 samples per brain region. RNA isolation was performed using the RNeasy^®^ Micro Kit (Qiagen) following the manufacturers instructions. 500 ng of each sample was converted into a sequencing-ready library using the CATS RNA-seq kit with rRNA depletion (Diagenode). The RNA integrity, effectiveness of the rRNA depletion and the final library sizes were verified by running aliquots on a TapeStation 2200 system with the appropriate ScreenTape devices (Agilent). Then, 12 samples for each brain region were pooled to equal amounts. The samples were sequenced on a total of 8 lanes using a HiSeq3000 instrument (Illumina) with a single end flowcell for 75 cycles. The obtained sequencing reads were demultiplexed with bcl2fastq version 2.20 with default parameters. Raw sequence data were preprocessed with a local Galaxy server (version 17.09), where FASTQ Groomer (version 1.1.1) and FASTQ Trimmer (version 1.1.1) were run (Blankenberg et al., 2010). The quality of the sequence data was checked by FastQC (version 0.11.8) (Andrews, 2010). The pseudoalignment with Kallisto (version 0.45.1) (Bray et al., 2016) was performed with the sequence data and an index built from the mouse transcriptome (Mus_musculus.GRCm38) with K-mer size = 21. The resulting Abundance.h5 files were imported to R (version 3.6.0) for differential gene expression analysis with the library DESeq2 (Love et al., 2014) and gene enrichment analysis with topGO (Alexa and Rahnenfuhrer, 2019). The detailed software setup and programming scripts can be found on Github. The data have been deposited in the SRA database under the accession number PRJNA628975.

### Cell culture, protein isolation for shRNA screening

NIH 3T3 mouse fibroblast cells (ATCCRCRL-1658) were maintained in Dulbecco’s modification of Eagle’s medium (DMEM, with 4.5 g/l glucose, L-glutamine & sodium pyruvate, Corning) with fetal bovine serum (FBS, 10% v/v, Biowest) and Penicillin/Streptomycin (100 U/mL, Roche Diagnostics GmbH). Cells were cultured on 10 cm sterile plates in 37 °C, in humidified chamber with 5% CO_2_. Lipofectamine 2000 (Invitrogen) was used to transfect cells (at 70% confluence, one day after seeding) according to manufacturer’s protocol. 10 μg of plasmid DNA (4 variants of shRNA carrying plasmids, OriGene TL501619) DNA was used to transfect one plate. The medium was replaced 6 hours after transfection with a standard one. 3 days after transfection cells were washed twice with cold PBS (137 mM NaCl, 7.97 mM Na_2_HPO_4_ × 12 H_2_O, 2.68 mM KCl, 1.47 mM KH_2_PO_4_), detached in PBS and moved to a 1.5 ml tube. After centrifugation the cell pellet was lysed in 1 ml of cold RIPA buffer (50 mM Tris-HCl pH7.4, 1% NP-40, 0.5% Na-deoxycholate, 0.1% SDS, 150 mM NaCl, 2 mM EDTA, 50 mM NaF) with freshly added protease (cOmplete ULTRA tablets, EDTA-free, Roche Diagnostics GmbH) and phosphatase inhibitors (PMSF 1mM, Na3VO4 0.2mM). The protein isolate was separated from cell debris by centrifugation (15 min, 14000g, 4°C) and stored at −20 °C.

### Protein isolation from brain tissue

Isolated brain regions from 3 animals were lysed in 500 μl cold lysis buffer (50 mM Tris-HCl, 150 mM NaCl, 0,25% SDS, 0,25% Sodium Deoxycholate, 1 mM EDTA) with freshly added protease (cOmplete ULTRA tablets, EDTA-free, Roche Diagnostics GmbH) and phosphatase inhibitors (PMSF 1 mM, Na3VO4 0.2 mM). Protein isolates were separated from cell debris by centrifugation (15 min, 14000 g, 4 °C) and stored at −20 °C.

### Western Blot

Protein sample concentration was determined by BCA (Pierce™ Rapid Gold BCA Protein Assay Kit). Samples were mixed with Laemmli buffer, incubated for 5 min at 95 °C and 80 μg of protein was separated on a 10% SDS-PAGE gel (alongside PageRuller™ Plus Prestained protein Ladder, Thermo Scientific). Samples were transferred onto a nitrocellulose membrane (Amersham Protan 0.45 μm, GE healthcare). Blocking was performed at 25 °C for 1 hour (5% milk, 1xTBS, 0,1% Tween), followed by 16 hours primary antibody incubation at 4 °C (rabbit anti-PER1 antibody produced by J. Ripperger, 1:1000; mouse anti-HSP90, 1:1000 (Santa Cruz Biotechnology, sc-13119); in 5% milk, 1xTBS, 0,1% Tween). Next, the membrane was washed 3 times with 1xTBS/0,1% Tween, incubated for 3 hours at 25 °C with the secondary antibody (anti-rabbit IgG peroxidase 1:10000, Sigma-Aldrich; anti-mouse IgG peroxidase 1:10000, Sigma-Aldrich; in 5% milk, 1xTBS,0,1% Tween) and washed 3 times with 1xTBS/0.1% Tween. After 5 min incubation with the developing reagent was added (Pierce^®^ ECL Western Blotting Substrate, 32106, Thermo Scientific) and images were captured using Azure 300 imaging system (Azure Biosystems).

### *In situ* hybridization

Animals were sacrificed at the experiment specific zeitgeber time. Tissue preparation, sectioning, α^35^S-UTP labeled riboprobe production and hybridization was followed according to (Albrecht et al., 1998).

*mPer1* and *mPer2* probes were produced from pBluescript SK(-) and pCR™II-TOPO^®^ respectively. The specificity of the antisense probe was verified using sense probe for hybridization. The signal obtained from the X-ray film (Amersham Hyperfilm MP, GE Healthcare) was quantified using densitometric analysis (GS-800, BioRad with Quantity One software, Biorad). In the case of weak signal, quantification was performed using silver-stained dark-field microscopy pictures (Zeiss Axioplan) coupled with Hoechst staining for nuclei. For both methods, the relative probe signal was calculated by subtracting the background signal taken from the neighboring region.

### Statistical analysis

Depending on experimental design, an appropriate method of statistical analysis was used (two-way ANOVA, one-way ANOVA, student t-test). Those statistics were calculated using GraphPad Prism software version 7.0d. A p-value lower than 0.05 was considered statistically significant. Circ Wave 1.4 (Hut, 2013) was used for cosinor analysis of expression patterns.

## Acknowledgments

We thank Antoinette Hayoz, Stéphanie Aebischer, Jean-Charles Paterna (Viral Vector Facility, University of Zürich) and the Bioimage platform (University of Fribourg) for technical support. Funding from the Velux Foundation (Projects 995 and 772) and the Swiss National Science Foundation (310030_184667/1) are acknowledged.

## Supplemental Figures

**Supplemental Figure 1**.

(**A**) Immobility time in the forced swim test (FST) of wild type male mice assessed over several days at ZT6 after no light pulse (LP) (black bars), after a LP at ZT14 (red bars), and after a LP at ZT22 (blue bars). Two-way RM ANOVA (n=13-15), ZT14 LP p=0.44, ZT22 LP p=0.14, values are means ± SEM. (**B**) Sucrose preference was tested allowing mice to choose between water or sucrose (1-5% (weight/vol)) over 3 days for each sucrose concentration. Two-way RM ANOVA revealed no differences between wild type (WT, n=21, black lines) and Per1^-/-^ mice (n=18, grey lines), p=0.67, values are means ± SEM.

**Supplemental Figure 2**.

(**A**) Light induction of Per1 at ZT14 in the LHb and the SCN. Dark-field images of coronal sections containing the habenula (rostral to caudal, left panels) and the SCN as positive control (right panels). The yellow signal represents the hybridization signal detecting Per1 mRNA and blue represents Hoechst-dye stained cell nuclei. The MHb can be distinguished from the LHb by the densely packed blue-colored nuclei. Brain section of mice sacrificed 60 min. after the light pulse (bottom row) and the corresponding controls are shown (top row). Scale bar: 200μm. (**B**) Brain region specific markers for verification of isolated brain regions used for further analysis. Quantitative PCR comparing various genes in the SCN, LHb, VTA and NAc. The most specific gene for the SCN is irs4, for the LHb it is gpr151, for the VTA it is tacr3 and for the NAc it is tac1. One-way ANOVA with Tukey’s multiple comparisons test was used, n=12-16, ****p<0.0001, values are means ± SEM.

**Supplemental Figure 3.**

(**A**) Characterization of shRNA against *Per1*. Western blot (left panel) using extracts from NIH 3T3 cells transfected with different shRNAs against *Per1*. The quantification of 3 experiments (right panel) revealed the best silencing activity for shRNA A and shRNA B. For further experiments shRNA A was chosen, because shRNA B had some similarities with *Per2*. (**B**) Brain sections of the habenular region are shown. Optimization of infection efficiency was performed by testing different variants of adeno-associated virus (AAV) expressing GFP (green color). Blue indicates cell nuclei (DAPI staining). For the lateral habenula (hatched white lines) AAV6 appeared to have the most localized and strongest infection potential. Scale bar: 200 μm.

**Supplemental Figure 4**.

Brain region-specific markers for verification of isolated brain regions used for further analysis. Quantitative PCR comparing various genes in the SCN, LHb, VTA and NAc. The most specific gene for the SCN is irs4, for the LHb it is gpr151, for the VTA it is tacr3 and for the NAc it is tac1.

