## Supplementary figures and images for "Light affects behavioral despair involving the clock gene *Period 1*"

### Supplemental Figure 1

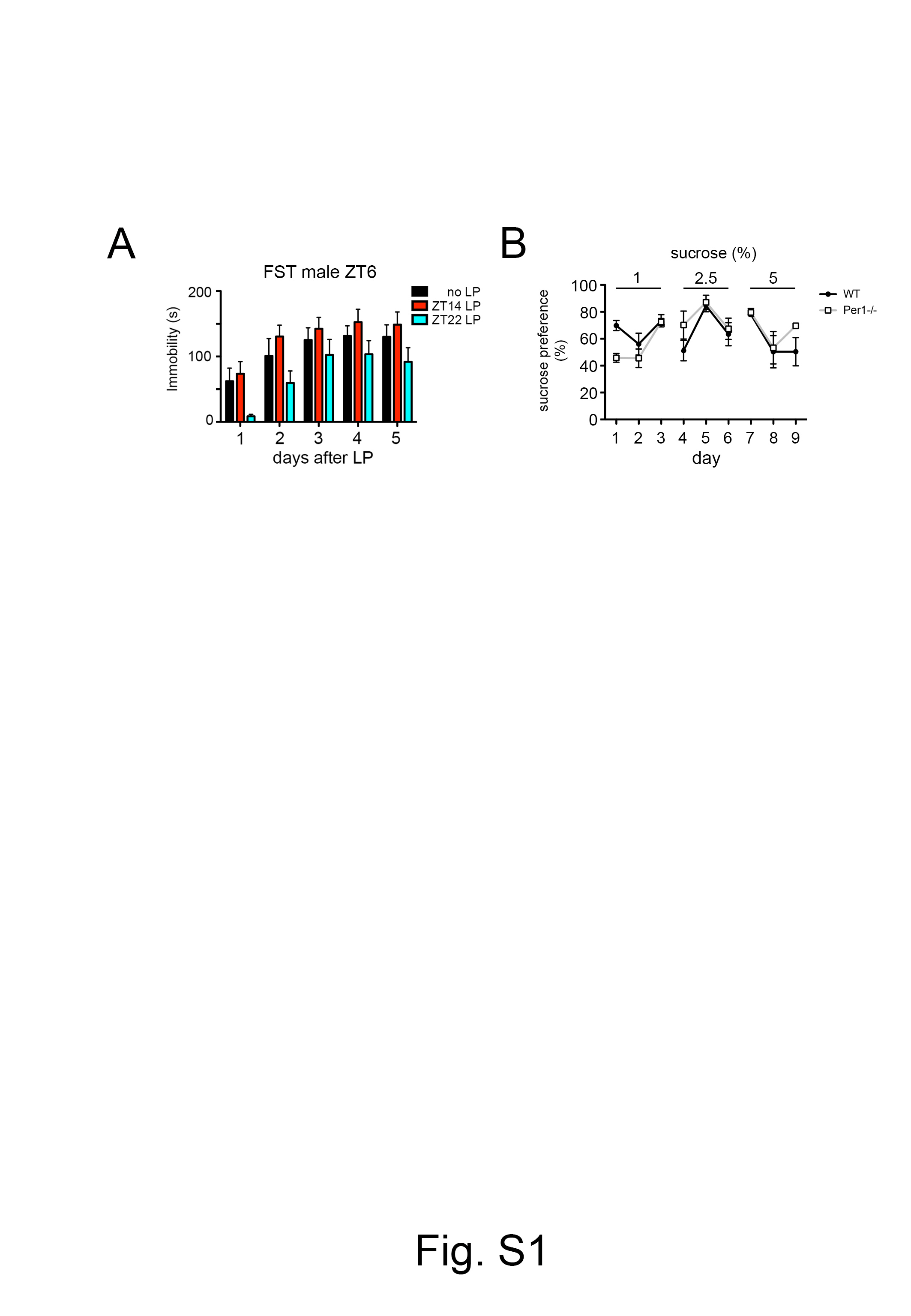

### Supplemental Figure 2

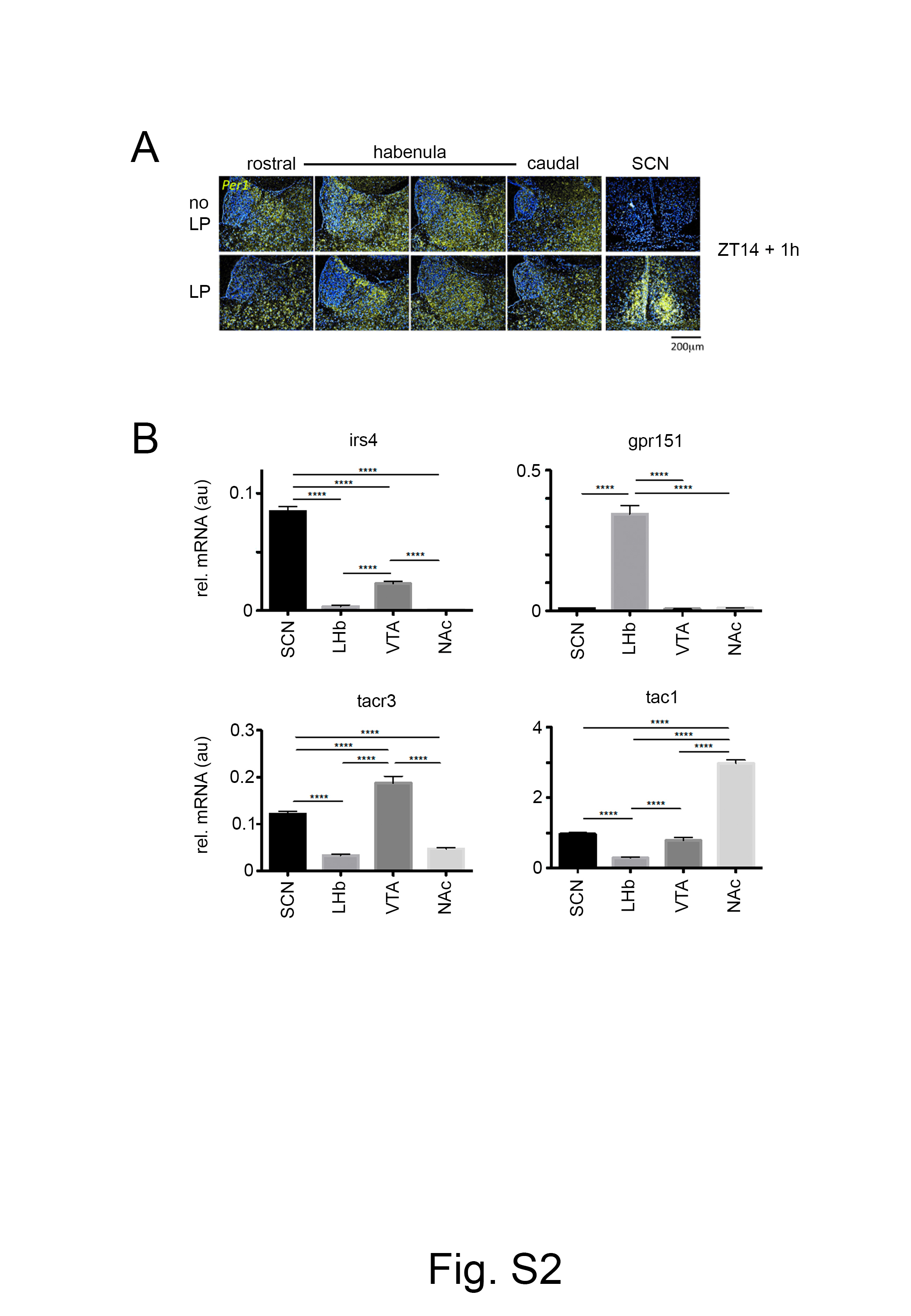

### Supplemental Figure 3

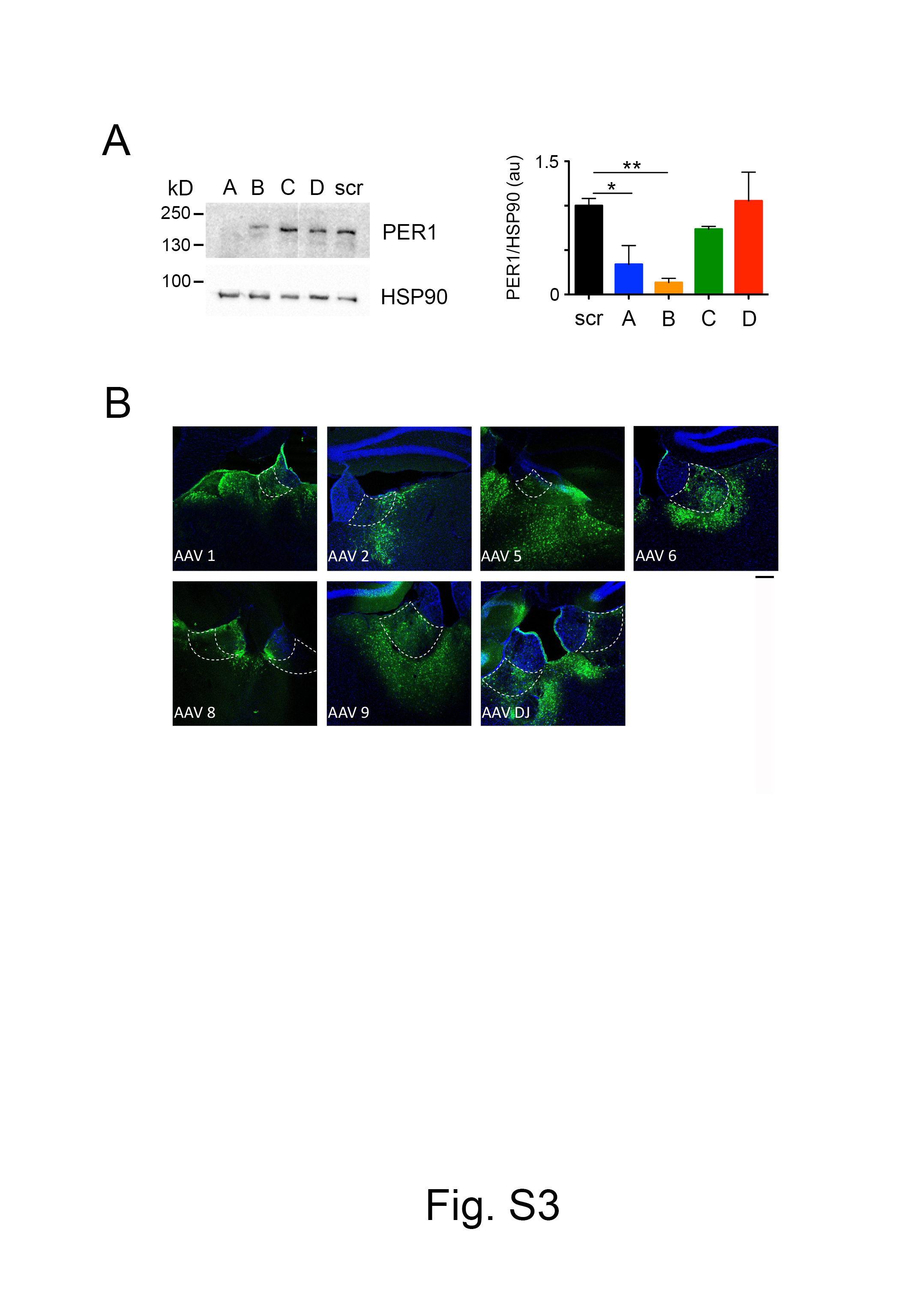

### Supplemental Figure 4

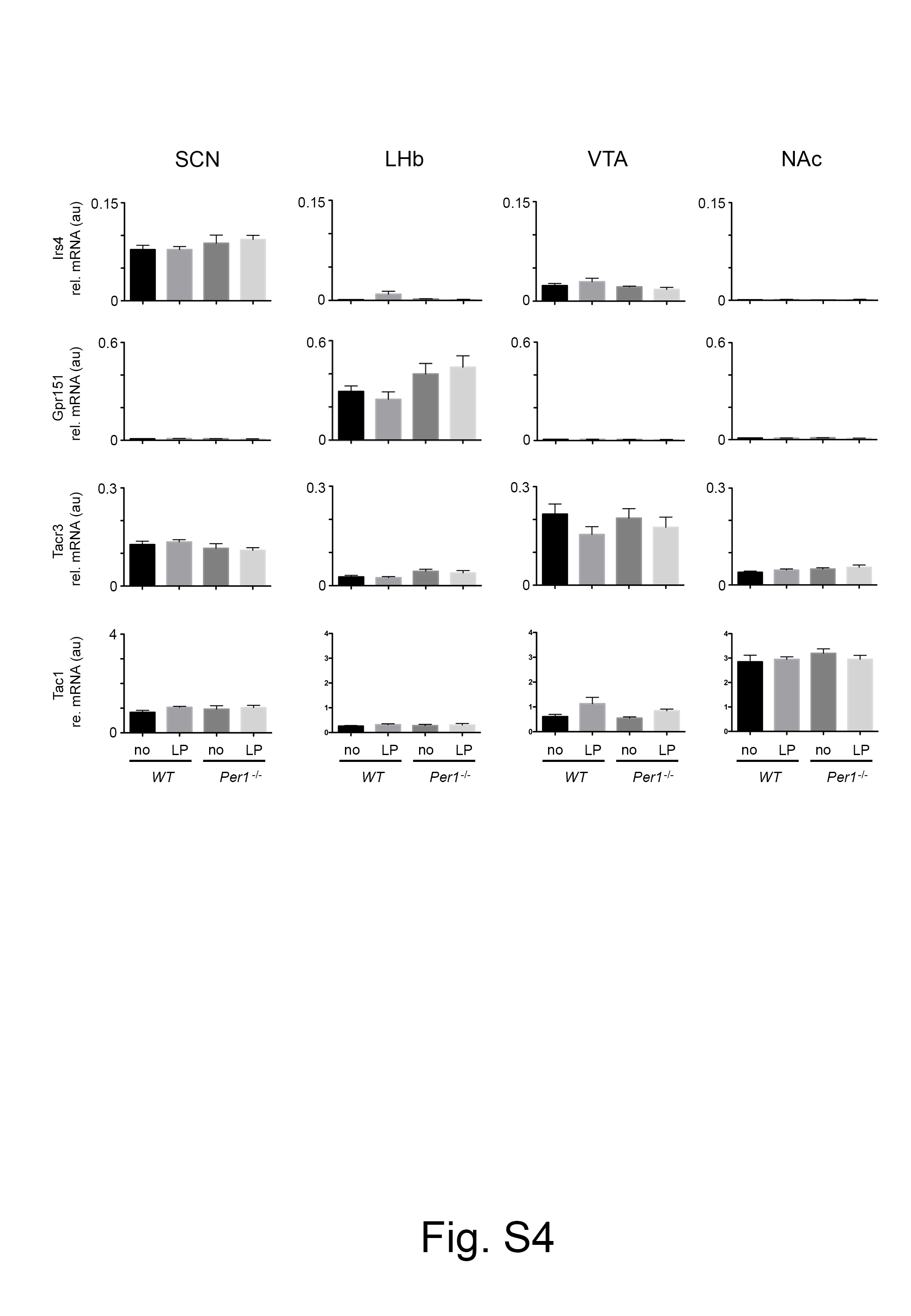
