## Supplementary material for "Light affects behavioral despair involving the clock gene *Period 1*": Table 10

| Gene | Primer/probe | Sequence |
| --- | --- | --- |
| *mPer1* | forward primer | GCC CCG CCT CCT TGC TAC A |
|  | reverse primer | ACT GGG GCC ACC TCC AGT TC |
|  | probe | TCC TTC CCT GCC AGT CCC CAA ACC CC |
| *mPer2* | forward primer | TCC ACA GCT ACA CCA CCC CTT A |
|  | reverse primer | TTT CTC CTC CAT GCA CTC CTG A |
|  | probe | CCG CTG CAC ACA CTC CAG GGC G |
| *mDrd3* | forward primer | CCT GGC TTC CCT CAG CAG TCT |
|  | reverse primer | GCT CCA TTT GTC CCG TGG CAT CT |
|  | probe | TGT CTG CGG CTG CAT CCC ATT CGG CA |
| *mComt* | forward primer | GTG GCT ACT CAG CCG TGC GA |
|  | reverse primer | GCT GGG TGA TGG CAG CGT AGT |
|  | probe | TGG CCC GCC TGC TGC CAC CT |
| *egfp* | forward primer | CAT CTG CAC CAC CGG CAA GC |
|  | reverse primer | GGT CGG GGT AGC GGC TGA A |
|  | probe | TGC CCG TGC CCT GGC CCA CC |
| *mGpr88* | forward primer | ACC AGC TAA GGA CTG GCC CC |
|  | reverse primer | GCC CTA GCG TTG GCT TCG TG |
|  | probe | CAG GCA AGT GCC AGG ATG GGC CAC GC |
| *m18SrRNA* | forward primer | CCG GCG GCT TGG TGA CTC TA |
|  | reverse primer | GGC AGA CGT TCG AAT GGG TCG T |
|  | probe | CCT CGG GCC GAT CGC ACG CC |
| *mTac1* | forward primer | AGA GCA AAG AGC GCC CAG CA |
|  | reverse primer | CGC CAC GGC CAC GAG GAT TT |
|  | probe | CCT GCG GAG CAT CCC CGC GG |
| *mTacr3* | forward primer | GGC TAA ACG AAA GGT TGT AA |
|  | reverse primer | AGA TCG CAG TGA GAA TGA AA |
|  | probe | GAC ATT TGC CAT CTG CTG GC |
| *mGrp151* | forward primer | CTG GCG GGC ATC TGG GTT GT |
|  | reverse primer | GCG TGA CGT CTG GTG GTG CTA A |
|  | probe | AGC CAG CCT GCT TCC CCT GCC AG |
| *mTspo* | forward primer | GGT CAG CTG GCT CTG AAC TG |
|  | reverse primer | CAG TCG CCA CCC CAC TGA CA |
|  | probe | TGC CCG GCA GAT GGG CTG GGC |
| *mTprkb* | forward primer | GGC TGG CAT CAG ACC CAC AGA |
|  | reverse primer | GGG CCC GTA GAG TCG GGA AA |
|  | probe | CCT GCG TCT GCC CTC TGA GGG CTG |
| *mIrs4* | forward primer | GGA CTT TGC CAG ACG AGA CT |
|  | reverse primer | TGG TTT TGG TGG CAG TGT AA |
|  | probe | CAG GAA GTG ACC ACT CTC CGA AAT GA |
| mRev-erbα | forward primer |  |
|  | reverse primer |  |
|  | probe |  |

**Table 1** Real-time PCR primers and probe sequences
